# fastlin: an ultra-fast program for *Mycobacterium tuberculosis* complex lineage typing

**DOI:** 10.1101/2023.07.11.548517

**Authors:** Romain Derelle, John Lees, Jody Phelan, Ajit Lalvani, Nimalan Arinaminpathy, Leonid Chindelevitch

## Abstract

Lineage typing of the *Mycobacterium tuberculosis* complex (MTBC) has evolved from traditional phenotypic methods to advanced molecular and genomic techniques. In this study we present fastlin, a bioinformatics tool designed for rapid MTBC lineage typing. Fastlin utilises an ultra-fast alignment-free approach to detect previously identified barcode single nucleotide polymorphisms (SNPs) associated with specific MTBC lineages directly from fastq files. In a comprehensive benchmarking against existing tools, fastlin demonstrated high accuracy and significantly faster running times. Analysis of large MTBC datasets revealed fastlin’s capability not only to predict MTBC lineages, but also to detect mixed-lineage strain mixtures and estimate their proportions. Fastlin offers a user-friendly and efficient solution for MTBC lineage typing, complementing existing tools and facilitating large-scale analysis.

## Introduction

Lineage typing of *Mycobacterium tuberculosis* complex (MTBC), which notably includes the aetiological agent of tuberculosis disease, has evolved from traditional phenotypic methods to more advanced molecular techniques. Early methods relied on observable characteristics in the laboratory and broad classifications, while DNA-based methods like RFLP and spoligotyping provided increased resolution but had technical limitations (Jagielski *et al*, 2014).

The advent of whole-genome sequencing (WGS) has revolutionised the comparative analysis of MTBC samples, allowing for high-resolution classification (Diel *et al*, 2019; Wyllie *et al*, 2018). While large genomic deletions were initially proposed as lineage markers, the use of single nucleotide polymorphisms (SNPs) as barcode markers has gained prominence due to their higher abundance, enabling more precise identification and classification of MTBC lineages (Cancino-Muñoz *et al*, 2019; Coll *et al*, 2014; Napier *et al*, 2020).

The identification of barcode SNPs traditionally involves the alignment of WGS reads to a reference genome, the basis of popular tools such as TB-profiler (Phelan *et al*, 2019) and TbLG (Shitikov and Bespiatykh 2023). As it is primarily designed for general variant calling, read alignment is a complex and computationally intensive approach that scales poorly to large datasets. To overcome this challenge, we propose a new bioinformatics tool, fastlin, which uses an alignment-free approach and performs MTBC lineage typing in seconds rather than minutes.

## Methods

Fastlin is written in rust and takes as input a SNP barcode file and the directory containing the fastq files to be typed. It is freely available at https://github.com/rderelle/fastlin and can easily be installed using Conda. The program essentially exploits the split-k-mer approach developed in SKA (Harris 2018), which uses k-mers split over a variable middle base, but with a significant speedup for lineage typing relying on the facts that: (i) fastlin does not need to compare and store all k-mers, only those relevant to barcode SNPs, and (ii) it typically does not need to scan all sequencing reads to identify barcode SNPs, assuming that the relevant reads are randomly distributed within the input fastq files.

The input barcode file contains all barcode SNPs with corresponding lineage names, and the 50 nucleotides upstream and downstream of each SNP, extracted from the genome of *M. tuberculosis* H37Rv; NC_000962.3. Fastlin then builds the barcode k-mers at the start of each run by combining (k-1)/2 downstream nucleotides, the barcode SNP and (k-1)/2 upstream nucleotides, with the k-mer size k being an odd number between 11 and 99 defined by the user (25 by default; see the Results section). The barcode k-mers are saved in memory together with their reverse complements. The input barcode file also contains the expected genome size, allowing fastlin to estimate the number of extracted k-mers needed to reach the user-defined k-mer coverage threshold (turned off by default).

K-mers are then extracted from the input fastq files and compared to the barcode k-mers until the entire fastq file is scanned, or until the number of extracted k-mers reaches the k-mer coverage threshold if one has been specified by the user. Once this process is finished, only those k-mers with a minimum number of occurrences are retained, and a lineage is validated if the number of retained k-mer barcodes reaches a user-defined threshold (3 by default). Finally, the coarser lineage names are removed (e.g., ‘4.1’ is removed if ‘4.1.2’ is detected), leading to the final lineage determination. If more than one lineage is detected, their abundances are calculated using the median of their barcode SNP occurrences. The relative frequency of each lineage can then be calculated as the ratio of their abundance divided by the sum of all final lineage abundances.

The barcode file used in this study and the Python scripts used to build and test it are available at https://github.com/rderelle/barcodes-fastlin. Sample analyses were performed using TB-profiler v5.0.0, QuantTB v1.01 and fastlin v0.1.0. Runtime benchmarks were performed with SKA v1.0 and TB-profiler v4.4.2 (with 6G RAM memory) and fastlin v0.1.0 on a Macbook Air laptop (M2, 8G RAM, 2022) using a single CPU. The 3139 MTBC samples were selected to represent all MTBC lineages as predicted by the TB-profiler database. The two most important contributors to this dataset were the Wellcome Trust and the UKHSA (formerly PHE), as part of the CRyPTiC consortium. All results mentioned in this manuscript, including runtimes, are available in Supplementary data 1.

## Results

For this case study we used the barcodes identified in TB-profiler (Phelan et al, 2019). At the time of writing, this set included 1101 SNPs defining 125 MTBC major and minor lineages, with a minimum of 4 and a maximum of 10 SNPs per lineage. Since these barcode SNPs were identified using read mapping, we first determined the minimal k-mer size at which fastlin did not generate false positive SNP calls. Using k-mer sizes ranging from 11 to 99 nucleotides and three high-quality genome assemblies with known lineages, we found that only one barcode SNP was incorrectly detected with k-mer sizes above 19 (Supplementary data 2). We discarded this problematic SNP barcode and set the default k-mer size to 25.

We then typed 3139 MTBC samples downloaded from the NCBI’s SRA database. To assess the quality of fastlin’s lineage predictions we compared them to the ones made by TB-profiler. Fastlin and TB-profiler predicted identical lineages for most samples, with disagreements found in only 68 samples (Supplementary data 3). Most of these disagreements were in putative mixed samples (n=51; 16 and 44 mixed samples identified by TB-profiler and fastlin respectively) or in samples for which either TB-profiler or fastlin failed to identify any lineage (8 and 1 samples, respectively). Only 2 pure samples resulted in lineage predictions in full disagreement.

We then re-analysed the 3139 MTBC samples with a maximum k-mer coverage threshold of 80x (the median k-mer coverage of all samples was 91x). Theoretically, with a minimum number of occurrences for each barcode SNP set to 4 by default, fastlin should be able to type all samples at this coverage threshold. Using lineage predictions obtained by fastlin with default parameters as a reference, we only found 5 samples with coarser (truncated) lineage predictions (e.g. predicted lineage ‘2.2.1’ instead of ‘2.2.1.2’; ERR3276004, ERR3276023 and SRR7453034) and 2 samples without a lineage prediction (ERR3276012 and ERR3275788), all corresponding to BAM-derived fastq files as suggested by the presence of a BAM file in the SRA database alongside the fastq files for these samples. Since BAM files are usually sorted by genomic position for further analyses, the assumption of a random read distribution in fastq files made by fastlin when a k-mer coverage threshold is set did not hold for these samples, and they consequently required more reads than expected to be correctly typed.

Next, we assessed fastlin’s ability to detect mixtures of strains by analysing a set of 47 pairs of *M. tuberculosis* samples initially produced to study cases of MTBC relapse and re-infection (Bryant *et al*, 2013) and subsequently re-analysed to test the MTBC mixed strain detection tool QuantTB (Anyansi *et al*, 2020). We ran QuantTB, TB-profiler and fastlin on this dataset. Eleven samples were identified as mixed by at least one of these three tools, with 9, 6, and 10 mixed samples detected by QuantTB, TB-profiler and fastlin respectively (Figure 1A and Supplementary data 4). The sample identified as a mixture of strains solely by QuantTB consists of a mixture of closely related strains (Bryant *et al*, 2013), possibly belonging to the same lineage (sample 8a; Supplementary data 4). Fastlin was the only tool to identify sample 45b as a mixed sample. We believe this sample indeed contains a strain mixture since (i) the lineage of its minor strain was also present in pure samples of this dataset and (ii) the minor strain was supported by 6 barcode SNPs. We repeated the fastlin analyses using maximum coverage thresholds ranging from 100x to 10x, in 10x increments. Fastlin successfully detected the 10 mixed samples down to a 60x maximum k-mer coverage (Supplementary data 4). Using simulated mixtures of pairs of strains from distinct lineages, we also found that fastlin provides reliable estimates of strain frequencies (RMSE = 2.47%; Supplementary data 4).

**Figure 1:**
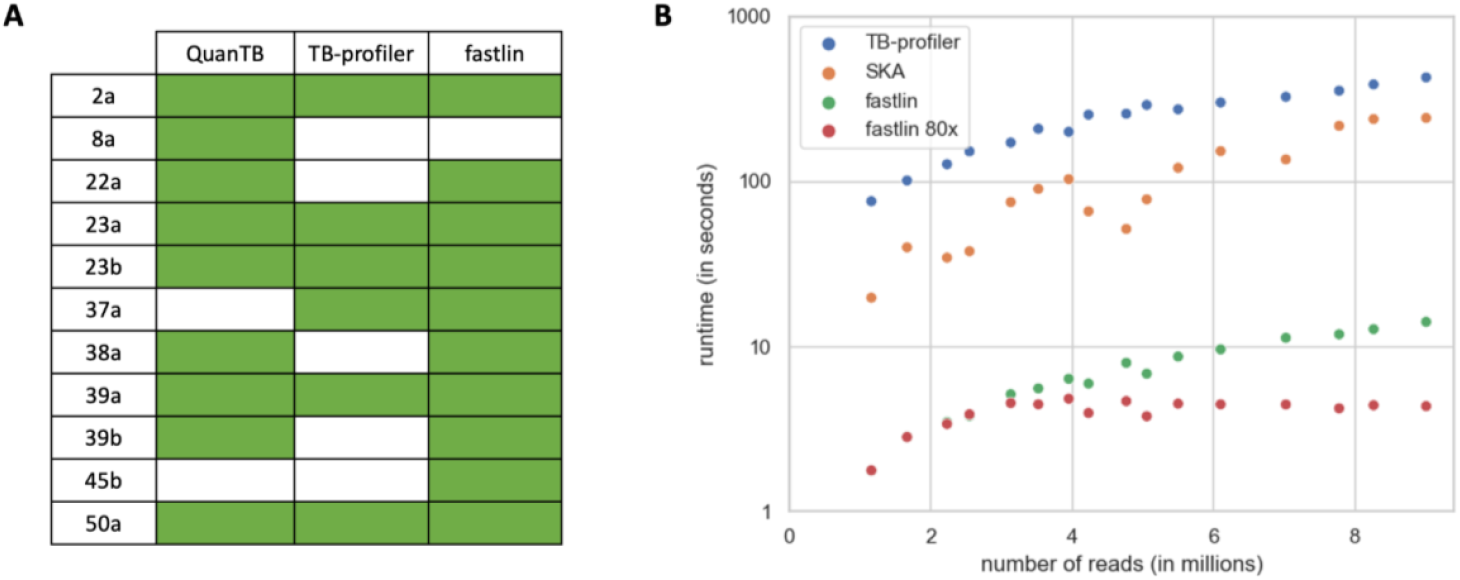
Fastlin benchmarking. **A**: Detected mixed samples in Bryant *et al* dataset. Samples with detected mixture are indicated with a green box. **B**: Runtimes obtained by TB-profiler, SKA (using k=27) and fastlin (with default parameters in green, and with a maximum k-mer coverage of 80x in red). Note the logarithmic scale on the y-axis.

Finally, we compared the running times of fastlin with those of SKA and TB-profiler, chosen as representatives of k-mer-based and read-mapping tools, respectively. It is important to note that these two bioinformatics tools are not fully dedicated to lineage typing and generate more information than fastlin does (e.g., all SNPs, drug resistance predictions). For this benchmark we used 16 SRA samples representing a large range of sequencing depths, all composed of paired-end 150bp reads. We observed that fastlin was at least an order of magnitude faster than SKA and TB-profiler, with running times ranging from 2 to 14 seconds per sample under default settings (Figure 1B). Using a maximum k-mer coverage threshold of 80x, fastlin’s running times were consistently under 5 seconds. Fastlin also exhibited minimal memory footprint, reaching a maximum use of only 4 MB in this benchmark, whereas both SKA and TB-profiler require several GB.

## Discussion

While primarily developed to achieve ultra-fast lineage typing, fastlin also shows accurate MTBC lineage prediction, comparable and in some cases superior to that of standard tools. Its lineage predictions are nearly identical to those made by TB-profiler and it shows an accurate identification of mixed-lineage samples and estimation of the associated lineage proportions. Fastlin also offers the added advantage of easy installation and use, eliminating the need for external programs and libraries required by mapping-based lineage typing methods. Thanks to its simplicity and efficiency, fastlin is an effective solution for rapidly typing large MTBC datasets.

Our analyses also uncover two limitations of fastlin. First, the presence of sorted reads in BAM-derived fastq files restricts the use of the maximum coverage threshold implemented in fastlin. Although this issue only affects the lineage prediction of seven files out of 3139 at a maximum coverage of 80x, this option is disabled by default, enabling fastlin to process all types of fastq files. However, in the case of in-house fastq files or fastq files of known provenance, we recommend specifying a maximum coverage threshold to further reduce fastlin’s runtime. Second, the detection of strain mixtures is intrinsically limited to the lineages defined by the SNP barcodes, and fastlin will fail to identify any mixture of strains belonging to the same lineage. If the identification of mixed samples is essential, we recommend the use of alternative tools with either larger reference datasets such as QuantTB (Anyansi *et al*, 2020) or database-free tools such as SplitStrains (Gabbassov *et al*, 2021).

While we tested fastlin with one specific set of MTBC barcode SNPs, new lineage classifications and their associated barcode SNPs have recently been proposed (Netikul *et al*, 2022; Shuaib *et al*, 2022; Coscolla *et al*, 2021; Shitikov and Bespiatykh 2023). To accommodate the evolving landscape of MTBC lineage typing, we made the scripts required to build and test custom barcode files available on GitHub. This will enable end-users to leverage fastlin with their barcode SNPs of choice. We recommend including a minimum of 4-5 SNPs per lineage, and considering all mutations rather than only synonymous SNPs, to ensure robust lineage classification.

## Supporting information

supplementary daya 1

supplementary daya 2

supplementary daya 3

supplementary daya 4

## Acknowledgments

This study was funded by the NIHR Health Protection Research Unit in Respiratory Infections, in partnership with the UK Health Security Agency (NIHR200927). The views expressed are those of the author(s) and not necessarily those of the NIHR or the Department of Health and Social Care. LC and NA acknowledge funding from the MRC Centre for Global Infectious Disease Analysis (reference MR/R015600/1), jointly funded by the UK Medical Research Council (MRC) and the UK Foreign, Commonwealth & Development Office (FCDO), under the MRC/FCDO Concordat agreement. For the purpose of open access, the authors have applied a Creative Commons Attribution (CC BY) licence to any Author Accepted Manuscript version arising.

