## supplementary daya 2 for "fastlin: an ultra-fast program for *Mycobacterium tuberculosis* complex lineage typing"

### Supplementary material 2

This study uses high-quality MTB genome assemblies downloaded from the Refseq NCBI database and infers their lineage as follows:

- 150bp single-end Illumina reads representing 60x coverage are simulated using Art-Illumina (Huang et al. 2012)
- the simulated reads are then typed using TB-profiler v4.4.2

The following table indicates the number of false positive k-mer barcodes (i.e. barcodes that should not be detected given the lineages of these genome assemblies) obtained at different k-mer sizes. K-mer sizes ranging from 61 to 99 are not shown (0 false positives in all cases). The false positive found with all three genomes at k-mer sizes ranging from 19 to 47 is always the same one: a G at position 1882572 (specific to the lineage 4.9.1).

| Kmer size | NC_000962.3 (lineage 4.9) | NZ_CP041872.1 (lineage 2.2.1) | NZ_CP041871.1 (lineage 3) |
|---|---|---|---|
| 11 | 871 | 860 | 875 |
| 13 | 359 | 353 | 363 |
| 15 | 62 | 63 | 62 |
| 17 | 14 | 14 | 14 |
| 19 | 1 | 1 | 1 |
| 21 | 1 | 1 | 1 |
| 23 | 1 | 1 | 1 |
| 25 | 1 | 1 | 1 |
| 27 | 1 | 1 | 1 |
| 29 | 1 | 1 | 1 |
| 31 | 1 | 1 | 1 |
| 33 | 1 | 1 | 1 |
| 35 | 1 | 1 | 1 |
| 37 | 1 | 1 | 1 |
| 39 | 1 | 1 | 1 |
| 41 | 1 | 1 | 1 |
| 43 | 1 | 1 | 1 |
| 45 | 1 | 1 | 1 |
| 47 | 1 | 1 | 1 |
| 49 | 0 | 0 | 0 |
| 51 | 0 | 0 | 0 |
| 53 | 0 | 0 | 0 |
| 55 | 0 | 0 | 0 |
| 57 | 0 | 0 | 0 |
| 59 | 0 | 0 | 0 |

Huang, W., L. Li, J. R. Myers, and G. T. Marth. 2012. 'ART: a next-generation sequencing read simulator', *Bioinformatics*, 28: 593-4.
