## supplementary daya 3 for "fastlin: an ultra-fast program for *Mycobacterium tuberculosis* complex lineage typing"

### Supplementary material 3

This file presents all 68 discrepancies between the lineage predictions made by TB-profiler and fastlin, and the 2 samples for which both TB-profiler and fastlin fail to infer any lineage.

#### Both TB-profiler and fastlin infer mixed samples but with different lineages (n=7)

| SRA ID | TB-profiler lineage | fastlin lineage | fastlin log barcodes |
|---|---|---|---|
| SRR11972381 | 4, 1.1.3 | 4.8 (9), 1.1.3 (7) | 1 (5, 7, 9, 7, 12, 5, 5, 5), 1.1 (6, 4, 6, 10, 12, 5, 7, 12), 1.1.3 (10, 7, 6, 12, 11, 7, 9, 4, 4), 4 (17, 10, 21, 8, 4, 16, 6, 20, 6, 15), 4.8 (8, 6, 11, 27, 11, 13, 9, 10, 8, 6) |
| SRR12199442 | 2, 1.2.2.2 | 1.2.2.2 (7), 1.2.1.2.1 (9) | 1 (17, 56, 21, 11, 10, 12, 4, 15, 16, 14), 1.1 (6), 1.1.3 (8), 1.2.1 (7, 10, 35, 14, 8, 11, 6, 8, 12), 1.2.1.2 (7, 6, 5, 14, 11, 10, 6, 69), 1.2.1.2.1 (17, 13, 5, 7, 7, 4, 21, 8, 10, 14), 1.2.2 (4, 5, 5, 74, 7), 1.2.2.2 (17, 7, 18, 6, 5), 2 (4), 2.2 (4, 8), 2.2.1 (34), 3.1.1 (4) |
| SRR19355499 | 4.2.2.2, 3.1.1, 2, 1 | 3.1.1 (6), 2.2.1 (6) | 1 (5), 1.1 (7), 2 (6, 4, 4, 7, 12, 9, 11, 11, 4, 4), 2.2 (4, 11, 4, 4, 10, 7, 13, 10), 2.2.1 (10, 5, 4, 6, 6, 4, 11), 3 (4, 7, 5), 3.1.1 (6, 4, 7, 9), 4.2 (4), 4.2.2 (4), 4.2.2.2 (4) |
| SRR19428521 | 2.2.1, 1.1.2 | 3.1.1 (11), 2.2 (5) | 2 (4), 2.2 (4, 6, 7, 4), 3 (5, 8, 8, 8, 5, 7, 15, 14), 3.1.1 (12, 9, 6, 12, 14, 10), 4.2 (4) |
| SRR998662 | 6.3.1, 4.1.3 | 4.1 (4), 4.6.2.2 (4), 6.3 (4) | 4 (5, 5, 12, 4, 7, 7, 11, 19, 7, 15), 4.1 (7, 4, 4, 4), 4.1.3 (4, 5), 4.6 (8, 7, 7, 4), 4.6.2 (6, 7, 7, 5, 7, 6), 4.6.2.2 (5, 4, 4, 4, 5, 4), 6 (4, 8), 6.3 (4, 4, 4, 5, 7), 6.3.1 (4, 4) |

#### TB-profiler inferred mixed samples but fastlin did not (n= 8)

| SRA ID | TB-profiler lineage | fastlin lineage | fastlin log barcodes |
|---|---|---|---|
| SRR11972253 | 2.2.1.1, 1.1.3 | 1.1.3 (12) | 1 (6, 9, 11, 4, 8, 8, 9), 1.1 (9, 8, 4, 9, 10, 7, 4, 4, 4), 1.1.3 (19, 5, 18, 4, 8, 13, 5, 12, 22), 2 (4), 2.2.1 (4), 2.2.1.1 (4) |
| ERR4797054 | 6.3, 2.1 | 2.2.1.2 (28) | 1.1.2 (74), 1.2.1.2.1 (47), 2 (34, 27, 22, 14, 22, 18, 22, 29, 28, 26), 2.2 (21, 38, 22, 26, 29, 19, 16, 25, 15, 25), 2.2.1 (14, 35, 37, 14, 7, 24, 20, 16, 39), 2.2.1.2 (36, 25, 31, 25) |
| SRR1181390 | 4.9, 2.2 | 2.2.1 (5) | 2 (7, 5), 2.2 (4, 6, 4), 2.2.1 (5, 8, 5, 5) |
| SRR1196677 | 2.2, La3 | 2.2.1 (6) | 1.1.2 (4), 1.2.1.2.1 (5), 2 (9, 5, 4, 4, 4, 5, 7), 2.2 (4, 4, 4, 5, 8), 2.2.1 (5, 7, 5, 6, 9) |
| SRR20969677 | 6.3, La3 | 2.1 (51) | 1.1.2 (193), 1.2.1.2.1 (185), 2 (49, 40, 47, 52, 55, 37, 35, 49, 94, 83), 2.1 (33, 49, 43, 73, 60, 95, 53, 91, 48, 38) |
| SRR023458 | 4.9, 4.1.2.1 | 4.1.2.1 (221) | 1.1 (6), 1.2.1 (4), 1.2.1.2.1 (5), 3.1.1 (8), 4 (342, 321, 442, 601, 401, 12, 299, 341, 392, 107), 4.1 (498, 582, 269, 572, 247, 438, 361, 528, 180, 52), 4.1.1.1 (5), 4.1.1.2 (4), 4.1.1.3 (5), 4.1.1.3.1 (4), 4.1.2 (142, 111, 277, 539, 271, 368, 136, 128, 392, 318), 4.1.2.1 (430, 161, 209, 216, 111, 315, 684, 467, 227, 168), 4.1.4 (4), 4.2.2.2 (5), 4.3.2 (4), 4.3.4.2.1 (4), 4.6 (29), 4.7 (6), 4.8 (4), 4.9 (4), 5.1.2 (4), 6.1.3 (4), La1.3 (7), La1.8.2 (4, 4) |
| ERR3283010 | 4.9;2.1, La1.8.1, La1.7.1 | La1.8.1 (4) | 8 (5), La1 (4, 5, 5), La1.8 (4, 5, 4, 4), La1.8.1 (4, 5, 4) |
| ERR4798262 | La2, La1.8.2, La1.2.BCG | 1.1.2 (16) | 1 (12, 19, 14, 18, 11, 14, 16, 24, 21, 20), 1.1 (13, 23, 20, 17, 23, 6, 11, 14, 23, 16), 1.1.2 (12, 17, 24, 16, 7, 16, 17, 18, 13, 19) |

#### fastlin inferred mixed samples but TB-profiler did not (n= 36)

| SRA ID | TB-profiler lineage | fastlin lineage | fastlin log barcodes |
|---|---|---|---|
| SRR19428508 | 1.1.2 | 4.1 (6), 1.1.2 (59) | 1 (48, 71, 55, 23, 55, 89, 66, 61, 99, 89), 1.1 (82, 84, 106, 89, 90, 59, 89, 73, 79, 94), 1.1.2 (29, 79, 62, 30, 59, 58, 59, 101, 95, 57), 4 (4, 4), 4.1 (5, 6, 7), 4.1.1 (6, 5), 4.1.1.1 (4, 5) |
| ERR176819 | 1.1.3.2 | 1.1.3.2 (72), 4.3.4.2 (4) | 1 (83, 72, 73, 95, 73, 66, 72, 70, 67, 83), 1.1 (99, 82, 77, 86, 58, 39, 82, 73, 82, 71), 1.1.3 (74, 50, 77, 68, 76, 79, 70, 70, 68, 79), 1.1.3.2 (58, 63, 66, 75, 79, 69, 75, 84), 2.2.1 (4), 4 (6), 4.2.2 (4), 4.3.4 (5, 5, 5), 4.3.4.2 (4, 4, 7), 4.3.4.2.1 (4) |
| ERR221621 | 1.1.3.2 | 1.2.2.2 (8), 1.1.3.2 (131) | 1 (148, 110, 148, 122, 135, 127, 148, 124, 129, 120), 1.1 (133, 160, 113, 125, 157, 120, 159, 125, 108, 100), 1.1.3 (134, 108, 111, 139, 147, 213, 114, 145, 125, 138), |

| | | | |
|---|---|---|---|
| | | | 1.1.3.2 (113, 148, 135, 143, 127, 101, 135, 104), 1.2.2 (6, 18, 9, 6, 7, 8, 8, 11, 8, 9), 1.2.2.2 (8, 5, 8, 7, 6, 8, 11, 6, 12) |
| ERR221666 | 1.1.3.2 | 3.1.1 (9), 1.1.3.2 (108) | 1 (125, 99, 118, 100, 103, 105, 124, 103, 93, 105), 1.1 (106, 123, 111, 129, 131, 92, 105, 102, 113, 122), 1.1.3 (102, 103, 132, 124, 116, 117, 123, 135, 119, 107), 1.1.3.2 (111, 82, 101, 106, 119, 115, 135, 98), 3 (11, 11, 6, 6, 4, 4, 10, 4, 6), 3.1.1 (12, 9, 9, 11, 9, 4, 6, 4) |
| SRR10809195 | 1.2.1 | 4.4.1.2 (5), 1.2.1 (104) | 1 (97, 145, 92, 113, 139, 51, 105, 109, 139, 120), 1.2.1 (125, 99, 96, 82, 109, 80, 123, 127, 129, 41), 4 (7, 4, 4, 4), 4.4 (7, 4), 4.4.1 (4, 5, 5, 4, 6), 4.4.1.2 (5, 5, 6, 4, 4) |
| SRR1011492 | 1.2.2.1 | 1.2.2.1 (93), 4 (4) | 1 (43, 104, 113, 12, 93, 120, 129, 85, 145, 42), 1.1.2 (4), 1.2.2 (145, 58, 116, 146, 54, 128, 120, 106, 83, 89), 1.2.2.1 (34, 41, 29, 32, 144, 124, 125, 91, 112, 95), 2.2 (4, 4), 4 (4, 5, 4), 4.3 (4, 4), 4.3.2 (5), 4.3.2.1 (5, 5) |
| SRR1013592 | 2 | 2 (106), 4 (4) | 2 (159, 113, 73, 92, 119, 116, 107, 101, 105, 21), 4 (5, 4, 4) |
| ERR4799973 | 3.1 | 3 (76), 1 (5) | 1 (5, 5, 7), 1.1 (4), 2 (4, 5), 2.2 (4), 2.2.1 (6, 6), 3 (53, 76, 63, 93, 94, 68, 79, 86, 76), 3.1 (78) |
| ERR688048 | 3 | 4.3.4.2 (6), 3 (384) | 1.2.1 (4), 3 (384, 416, 451, 395, 353, 219, 452, 360, 361), 4 (4, 4, 4, 6, 5), 4.1.4 (4), 4.3 (4, 4, 6, 4), 4.3.4 (4, 12, 4), 4.3.4.2 (6, 8, 5, 7, 6), 4.6 (4), 5.1.2 (6), 8 (4) |
| SRR4035524 | 3 | 3 (257), 4.1.2 (6) | 3 (205, 277, 308, 271, 161, 99, 257, 304, 236), 4 (4, 4), 4.1.2 (6, 4, 10), 4.2 (4) |
| ERR400377 | 4.1.2 | 4.1.2 (71), 4.3.4.2 (5) | 4 (89, 71, 70, 75, 50, 110, 70, 108, 26, 95), 4.1 (148, 96, 83, 104, 74, 76, 77, 58, 105, 72), 4.1.2 (72, 105, 70, 71, 55, 70, 81, 103, 65, 113), 4.3.4.2 (7, 5, 4), 4.3.4.2.1 (12) |
| ERR2514497 | 4.1.3 | 4.1.3 (98), 1.1.2 (5) | 1 (4), 1.1 (6, 8, 4), 1.1.2 (4, 7, 4, 7), 4 (108, 116, 92, 109, 110, 88, 119, 67, 131, 71), 4.1 (138, 114, 131, 120, 103, 106, 102, 78, 132, 97), 4.1.3 (108, 106, 83, 108, 115, 91, 75, 90, 141, 75) |
| SRR998798 | 4.1.3 | 4.6.2.2 (8), 4.1.3 (64) | 4 (69, 64, 62, 72, 82, 68, 62, 37, 84, 56), 4.1 (90, 78, 62, 79, 71, 74, 48, 53, 93, 70), 4.1.3 (67, 79, 44, 54, 83, 53, 62, 74, 61, 74), 4.6 (6, 6, 7), 4.6.2 (4, 5, 5, 6, 9, 7, 8, 5), 4.6.2.2 (8, 10, 4) |
| ERR234686 | 4.2.1 | 4.2.1 (112), 2.2.1 (4) | 2 (7, 4, 4, 6, 5), 2.2.1 (4, 4, 5), 4 (105, 118, 102, 110, 94, 139, 106, 123, 84, 122), 4.2 (113, 87, 104, 113, 114, 95, 87, 108, 98, 116), 4.2.1 (115, 96, 85, 126, 109, 121, 119, 102, 120, 106) |
| SRR20969577 | 4.2.2 | 4.2.2 (247), 2.2.1.1 (19) | 2 (21, 15, 30, 7, 20, 21, 14, 16, 18, 20), 2.2 (16, 16, 19, 22, 15, 27, 30, 7, 19, 25), 2.2.1 (27, 21, 11, 14, 13, 13, 15, 14, 22), 2.2.1.1 (16, 18, 21, 31, 25, 15), 4 (249, 256, 245, 240, 277, 225, 257, 256, 267, 221), 4.2 (242, 223, 267, 244, 285, 149, 287, 230, 259, 200), 4.2.2 (210, 258, 217, 254, 259, 191, 250, 244) |
| SRR5818596 | 4.2 | 3 (4), 4.2 (33) | 3 (4, 9, 4), 4 (45, 33, 37, 32, 46, 19, 48, 28, 23, 36), 4.2 (45, 35, 65, 73, 35, 12, 31, 31, 26, 25), 4.6 (32) |
| ERR8699189 | 4.3.2.1 | 2.2 (5), 4.3.2.1 (137), 3 (10) | 2.2 (6, 5, 5), 3 (12, 6, 11, 10, 13, 7, 6, 19, 9), 4 (171, 147, 174, 162, 119, 149, 116, 126, 126, 136), 4.3 (167, 159, 140, 126, 139, 96, 133, 166, 140), 4.3.2 (140, 145, 145, 148, 168, 105, 143, 139, 170, 168), 4.3.2.1 (126, 143, 134, 176, 140, 118, 156, 128, 118, 169) |
| ERR212171 | 4.3.4.2.1 | 4.3.4.2.1 (116), 3 (5) | 3 (5, 4, 5), 3.1.1 (6), 4 (153, 150, 117, 70, 164, 162, 117, 116, 89, 146), 4.3 (104, 145, 156, 112, 96, 110, 74, 155, 83), 4.3.2 (6), 4.3.4 (83, 89, 135, 144, 142, 92, 111), 4.3.4.2 (67, 97, 159, 151, 69, 162, 116, 72, 150, 129), 4.3.4.2.1 (85, 119, 140, 115, 126, 68, 118, 111) |
| SRR847802 | 4.4.1.1.1 | 4.9 (5), 4.1.2.1 (5), 4.4.1.1.1 (818), 2.2.1 (4) | 1.1 (5), 1.2.1.2 (4), 1.2.1.2.1 (13), 1.2.2.1 (8), 1.2.2.2 (5), 2 (6), 2.1 (4), 2.2.1 (4, 8, 4), 2.2.1.1 (7), 3 (4), 3.1.1 (5, 7), 3.1.2.2 (4), 3.1.3 (4), 4 (644, 864, 721, 837, 825, 740, 774, 731, 820, 791), 4.1 (4, 4, 5, 4, 4, 5, 7), 4.1.1.1 (5, 4), 4.1.1.2 (13), 4.1.2 (6, 4, 6, 4), 4.1.2.1 (4, 6, 5, 4, 9), 4.1.2.1.1 (4), 4.2 (4), 4.2.1.1 (4), 4.3.4 (4), 4.4 (844, 854, 767, 759, 883, 850, 851, 817, 819, 655), 4.4.1 (729, 789, 890, 823, 716, 845, 931, 772, 773, 769), 4.4.1.1 (915, 934, 807, 783, 958, 826, 798, 640, 859, 840), 4.4.1.1.1 (818, 985, 504, 821, 877, 759, 749, 911, 733), 4.4.2 (4), 4.6 (5), 4.6.1.2 (6), 4.6.3 (4), 4.6.5 (4), 4.7 (4), 4.8.1 (4), 4.9 (5, 5, 4), 4.9.1 (4, 4), 5 (8), 5.1.2 (23), 5.1.3 (5), 5.1.4 (5), 5.2 (6), 5.3 (4), 6.1.3 (4, 4), 7 (4, 5), 8 (11), 9 (5), La1.2.BCG (4), La1.3 (6, 7), La1.4 (4), La1.7.X-unk4 (8), La1.8 (4), La3 (9) |
| SRR4037763 | 4.4.1.1 | 2.2.1 (5), 4.4.1.1 (85) | 2 (4, 4, 8, 7, 5, 5), 2.2 (5, 13, 10, 5, 5, 7, 8, 7), 2.2.1 (9, 5, 5, 9, 6, 4), 4 (69, 56, 80, 69, 71, 72, 95, 77, 92, 78), 4.4 (85, 129, 61, 43, 112, 76, 69, 49, 133, 68), 4.4.1 (60, 73, 110, 78, 68, 87, 35, 30, 53, 71), 4.4.1.1 (52, 76, 125, 130, 91, 87, 52, 90, 75, 83), 8 (4) |
| SRR15368744 | 4.4.2 | 2.2 (6), 4.4.2 (250) | 2 (9, 7, 4, 4), 2.2 (6, 4, 7), 2.2.1 (4, 5), 2.2.1.1 (4), 4 (299, 223, 318, 275, 268, 225, 318, 309, 211, 275), 4.4 (268, 305, 298, 228, 227, 334, 317, 333, 295, 223), 4.4.2 (222, 272, 229, 307, 193, 284), 6 (4) |
| SRR8651637 | 4.4.2 | 2.2 (4), 4.4.2 (321) | 2.2 (4, 4, 4), 2.2.1 (4), 4 (252, 317, 365, 235, 197, 283, 329, 233, 353, 301), 4.4 (319, 130, 387, 303, 166, 451, 299, 288, 335, 365), 4.4.2 (316, 305, 326, 383, 350, 205) |
| ERR1199115 | 4.6.1.2 | 4.3.4.2.1 (7), 4.6.1.2 (123) | 4 (87, 122, 120, 31, 168, 160, 136, 94, 127, 115), 4.3 (4, 5, 7, 4, 7, 9, 7, 6), 4.3.4 (4, 6, 5, 5, 4), 4.3.4.2 (9, 9, 8, 6, 5, 5, 4), 4.3.4.2.1 (8, 5, 7, 8, 5, 8, 4), 4.6 (147, 118, 86, 123, 55, 114, 133), 4.6.1 (102, 109, 150, 78, 282, 93, 148, 116, 81, 121), 4.6.1.2 (130, 125, 121, 121, 164, 102, 122, 120, 130, 170) |
| ERR2707224 | 4.6.1.2 | 4.3.4.2.1 (7), 4.6.1.2 (123) | 4 (87, 122, 120, 31, 168, 160, 136, 94, 127, 115), 4.3 (4, 5, 7, 4, 7, 9, 7, 6), 4.3.4 (4, 6, 5, 5, 4), 4.3.4.2 (9, 9, 8, 6, 5, 5, 4), 4.3.4.2.1 (8, 5, 7, 8, 5, 8, 4), 4.6 (147, 118, 86, 123, 55, 114, 133), 4.6.1 (102, 109, 150, 78, 282, 93, 148, 116, 81, 121), 4.6.1.2 (130, 125, 121, 121, 164, 102, 122, 120, 130, 170) |
| SRR18213698 | 4.6 | 1.1.3 (4), 4.6 (66) | 1 (4, 6), 1.1 (4, 4), 1.1.3 (4, 4, 5), 4 (66, 40, 101, 45, 69, 39, 87, 66, 73, 82), 4.6 (81, 46, 67, 66, 65, 78, 64) |
| SRR2101048 | 4.6 | 1.1.3 (4), 4.6 (66) | 1 (4, 6), 1.1 (4, 4), 1.1.3 (4, 4, 5), 4 (66, 40, 101, 45, 69, 39, 87, 66, 73, 82), 4.6 (81, 46, 67, 66, 65, 78, 64) |
| ERR266580 | 4.8.2 | 4.8.2 (65), 4.6 (6) | 4 (59, 64, 33, 96, 43, 74, 68, 52, 74, 68), 4.6 (5, 6, 6), 4.6.3 (5), 4.8 (56, 63, 76, 59, 60, 23, 89, 69, 66, 52), 4.8.2 (55, 96, 66, 78, 70, 74, 54, 51, 33, 65), 5.1.2 (4) |

| | | | |
|---|---|---|---|
| ERR245719 | 4.9.1 | 4.3.4.2.1 (7), 4.9.1 (159) | 1.2.1.2.1 (5), 4 (155, 164, 268, 193, 163, 151, 196, 160, 253, 118), 4.1.4 (6), 4.2.2 (4), 4.3 (4, 4, 4, 7, 5, 6, 8, 7), 4.3.4 (8, 13, 4, 7), 4.3.4.2 (4, 7, 4, 5, 6, 4, 6, 5, 7), 4.3.4.2.1 (7, 7, 4, 7, 4), 4.6 (7), 4.9 (117, 221, 174, 193, 152, 186, 188, 201, 231, 181), 4.9.1 (165, 125, 158, 201, 159, 239, 227, 111, 130), La1.7.X-unk4 (6) |
| ERR4799905 | 4.9 | 4.9 (110), 2 (5) | 2 (4, 5, 8), 2.2 (5), 2.2.1 (7, 4), 3 (6), 4 (116, 84, 100, 92, 99, 90, 82, 100, 105, 108), 4.9 (86, 125, 125, 75, 101, 83, 119, 120, 123, 58) |
| ERR3801606 | 6.1.3 | 6.1.3 (228), 4.1.2.1 (6) | 4 (6, 12, 6, 4, 6, 8, 5), 4.1 (12, 7, 7, 4, 4, 7, 5), 4.1.2 (7, 5, 5, 5, 6), 4.1.2.1 (4, 4, 8, 9, 5, 10), 6 (236, 111, 172, 209, 196, 170, 179, 284, 181, 175), 6.1.3 (262, 242, 215, 191), La1.3 (5) |
| ERR9787253 | 6.1.3 | 6.1.3 (228), 4.1.2.1 (6) | 4 (6, 12, 6, 4, 6, 8, 5), 4.1 (12, 7, 7, 4, 4, 7, 5), 4.1.2 (7, 5, 5, 5, 6), 4.1.2.1 (4, 4, 8, 9, 5, 10), 6 (236, 111, 172, 209, 196, 170, 179, 284, 181, 175), 6.1.3 (262, 242, 215, 191), La1.3 (5) |
| SRR998606 | 6.2.1 | 6.2.1 (93), 6.3.1 (5) | 6 (103, 109, 58, 76, 103, 67, 73, 111, 116, 102), 6.2 (86, 104, 102, 78, 85, 112, 69, 94, 91), 6.2.1 (104, 83, 104, 69), 6.3 (4, 5, 9, 4, 4, 8), 6.3.1 (5, 9, 4, 4, 7, 6) |
| SRR998607 | 6.2.1 | 6.3 (4), 6.2.1 (87) | 6 (92, 71, 92, 78, 68, 90, 84, 129, 82, 87), 6.2 (77, 88, 101, 55, 66, 113, 74, 106, 70), 6.2.1 (125, 87, 88, 53), 6.3 (6, 4, 5, 4, 4, 4, 9), 6.3.1 (4, 7) |
| ERR3170439 | 6.3.3 | 6.3.3 (83), 4 (4) | 4 (4, 4, 4, 4), 4.6 (4), 4.6.2 (4), 4.6.2.2 (4), 6 (113, 100, 65, 99, 105, 113, 91, 113, 119, 109), 6.3 (107, 78, 89, 106, 91, 93, 79, 139, 85), 6.3.3 (83, 128, 83, 89, 81, 105, 77, 125, 84, 78) |
| SRR998576 | 6.3.3 | 6.3.3 (62), 6.3.1 (6) | 6 (84, 58, 61, 72, 74, 57, 76, 72, 61, 88), 6.3 (56, 64, 82, 76, 67, 101, 67, 64, 57), 6.3.1 (6, 4, 6), 6.3.3 (42, 62, 57, 54, 64, 55, 74, 63, 68, 84) |
| ERR234682 | La3 | 4 (5), La3 (97) | 4 (5, 5, 7, 5, 4, 4, 4), 4.8 (6, 4), La3 (112, 101, 91, 97, 97) |

#### TB-profiler failed to infer any lineage (n=8)

| SRA ID | TB-profiler lineage | fastlin lineage | fastlin log barcodes |
|---|---|---|---|
| ERR4819092 | | 1.2.1.2.1 | 1 (7, 4, 8, 5), 1.2.1 (11, 7, 5, 10), 1.2.1.2 (4, 5, 6, 5, 5), 1.2.1.2.1 (10, 4, 5, 6, 6, 9, 5, 5) |
| SRR21735256 | | 1.2.2.2 | 1 (4, 10, 13, 31, 10, 5), 1.2.2 (13, 8, 4, 45, 8, 7), 1.2.2.2 (7, 16, 11, 33), 4.1.2 (12) |
| SRR12199408 | | 4 (6), 1.2.2.2 (23) | 1 (24, 23, 36, 46, 35, 30, 17, 12, 11, 25), 1.1.1 (17), 1.2.2 (18, 36, 29, 10, 22, 35, 35, 18, 20, 10), 1.2.2.2 (28, 37, 15, 23, 37, 28, 18, 23, 23), 2.2 (5, 5), 4 (10, 6, 5), 4.3 (4) |
| ERR6358755 | | 4.6.1.1 | 1.1.2 (58), 1.2.1.2.1 (68), 2.1 (4), 4 (91, 47, 66, 65, 10, 112, 98, 51, 40, 69), 4.3.4 (4), 4.3.4.2 (6), 4.3.4.2.1 (4, 4), 4.4 (82), 4.6 (75, 50, 59, 55, 35, 66, 57), 4.6.1 (61, 74, 98, 38, 118, 54, 75, 79, 56, 68), 4.6.1.1 (60, 67, 50, 103, 64, 75, 63, 59, 76, 46) |
| SRR5341203 | | 3 | 3 (7, 9, 18, 17, 16, 6, 4, 17, 12) |
| ERR4187744 | | La1.8.1 | La1 (4, 6, 9), La1.8 (8, 7), La1.8.1 (13, 7, 5, 4) |
| ERR3468577 | | La1.8.1 | La1 (4), La1.8 (8, 5, 4), La1.8.1 (5, 6, 9, 6) |
| ERR4769482 | | La1 | La1 (5, 4, 4), La1.3 (5, 4) |

#### fastlin failed to infer any lineage (n=1)

| SRA ID | TB-profiler lineage | fastlin lineage | fastlin log barcodes |
|---|---|---|---|
| ERR025887 | 4.3.4.2, 1.2.1.2.1 | | 1 (4), 1.2.1 (6), 1.2.1.2 (4), 1.2.1.2.1 (4, 4), 4.3.4 (5) |

#### Both TB-profiler and fastlin failed to infer any lineage (n=2)

| SRA ID | TB-profiler lineage | fastlin lineage | fastlin log barcodes |
|---|---|---|---|
| ERR2747660 | | | La1 (6, 6), La1.8 (6, 5), La1.8.1 (10) |
| ERR4769545 | | | La1 (4) |

#### fastlin provided a more precise lineage prediction (n=6)

| SRA ID | TB-profiler lineage | fastlin lineage | fastlin log barcodes |
|---|---|---|---|
| SRR10305655 | 2.2.1 | 2.2.1.1 | 2 (244, 259, 264, 242, 225, 197, 198, 234, 245, 266), 2.2 (191, 250, 279, 187, 234, 207, 275, 256, 202, 220), 2.2.1 (224, 199, 270, 237, 203, 248, 239, 204, 269), 2.2.1.1 (9, 7, 5, 4, 7) |
| SRR1062937 | 4.6.1 | 4.6.1.1 | 4 (16, 22, 7, 6, 17, 9, 9, 9, 13), 4.6 (14, 9, 20, 15, 4, 5, 15), 4.6.1 (15, 7, 9, 11, 10, 9, 8, 9, 11, 10), 4.6.1.1 (17, 15, 20, 10, 8, 6, 7, 13) |
| ERR1213918 | 4.6 | 4.6.2.2 | 4 (10, 33, 16, 36, 37, 16, 29, 111, 29, 45), 4.6 (43, 62, 23, 46, 13, 40, 34), 4.6.2 (25, 15, 9, 41, 13, 48, 26, 93, 33, 41), 4.6.2.2 (7, 39, 57, 43, 37, 35, 8, 39, 20) |
| ERR9787030 | 6 | 6.1.2 | 6 (65, 105, 41, 60, 59, 82, 94, 96, 100, 100), 6.1.2 (60, 59, 120, 103, 81, 115) |
| ERR3468727 | La1.8 | La1.8.1 | La1 (4, 5), La1.8 (5, 5, 4, 6), La1.8.1 (5, 4, 5) |
| SRR10993935 | La1 | La1.6 | La1 (7, 5), La1.6 (5, 5, 5) |

#### TB-profiler and fastlin lineage predictions totally disagree (n=2)

| SRA ID | TB-profiler lineage | fastlin lineage | fastlin log barcodes |
|---|---|---|---|
| ERR9787014 | La2 | 6.1.1 | 4.1.1.1 (63), 6 (136, 86, 148, 115, 118, 90, 126, 104, 122, 155), 6.1.1 (122, 128, 118, 124, 149, 129, 150, 157, 146, 122) |
| ERR9787139 | 2.1 | 6.3.1 | 1.1.2 (73), 6 (119, 58, 60, 106, 101, 80, 104, 123, 123, 87), 6.3 (96, 84, 116, 55, 95, 161, 72, 120, 103), 6.3.1 (109, 92, 57, 82, 78, 102, 88, 130, 88, 91) |
