## supplementary daya 4 for "fastlin: an ultra-fast program for *Mycobacterium tuberculosis* complex lineage typing"

### Supplementary material 4

Same table as Figure 1a but with the relative frequencies of the minor strains indicated when detected (sample 23a had been sequenced twice):

| | QuanTB | TBprofiler | fastlin |
|---|---|---|---|
| 2a | 0.18 | 0.16 | 0.16 |
| 8a | 0.47 | | |
| 22a | 0.30 | | 0.07 |
| 23a | 0.33 / 0.40 | 0.31 / 0.38 | 0.34 / 0.28 |
| 23b | 0.17 | 0.15 | 0.15 |
| 37a | | 0.06 | 0.07 |
| 38a | 0.10 | | 0.07 |
| 39a | 0.27 | 0.19 | 0.28 |
| 39b | 0.06 | | 0.07 |
| 45b | | | 0.06 |
| 50a | 0.25 | 0.30 | 0.33 |

Regarding the mixed sample 8a identified solely by QuantTB, it should be noted that both TB-profiler and fastlin identified the major strain with the correct frequency value of 0.6, suggesting that a second strain was present in this sample (see Supplementary material 1). Upon close examination of the barcode SNPs identified by fastlin and their occurrences, we can hypothesise that the 'missing' minor strain belongs to lineage 4.4.1.1 but not 4.4.1.1.1 or any sub-lineage defined by the set of barcode SNPs.

Mixed samples identified by fastlin in the Bryant *et al* dataset at different maximum coverage thresholds ('X' indicates that a mixture of strains was detected by fastlin):

| samples | 10x | 20x | 30x | 40x | 50x | 60x | 70x | 80x | 90x | 100x |
|---|---|---|---|---|---|---|---|---|---|---|
| 2a | | X | X | X | X | X | X | X | X | X |
| 22a | | | | | X | X | X | X | X | X |
| 23a | X | X | X | X | X | X | X | X | X | X |
| 23b | | X | X | X | X | X | X | X | X | X |
| 37a | | | | | | X | X | X | X | X |
| 38a | | | | | X | X | X | X | X | X |
| 39a | | | | | | X | X | X | X | X |
| 39b | | | X | X | X | X | X | X | X | X |
| 45b | | | | | X | X | X | X | X | X |
| 50a | X | X | X | X | X | X | X | X | X | X |

We assessed the accuracy of minor strain frequency estimations made by fastlin in cases of mixed samples. To this end, we simulated Illumina reads representing mixed samples with minor strain frequencies ranging from 0.1 to 0.5 in increments of 0.1 (each sample with a total coverage of 60X (minor + major strain); 5 replicates at each frequency). We used the ART-illumina v2.5.8 tool (Huang et al. 2012) to generate 150bp single-end reads from two genome assemblies with known lineages (see Supplementary data 1): H37Rv genome assembly for the major strain (NC_000962.3; lineage 4.9) and genome assembly NZ_CP041804.1 for the minor strain (lineage 4.8).

With default parameters, fastlin correctly identifies all 25 simulated samples as mixtures of strains. The following plot shows the linear regression of the minor strain frequencies inferred by fastlin (95% confidence interval in light blue; $R^2$= 0.97; p-value= 5.55 x $10^{-19}$). We then measured the deviation of all estimates from their true values using the Root Mean Square Error (RMSE) and obtained a value of 2.47%, indicating that fastlin provides reliable estimates of strain frequencies.

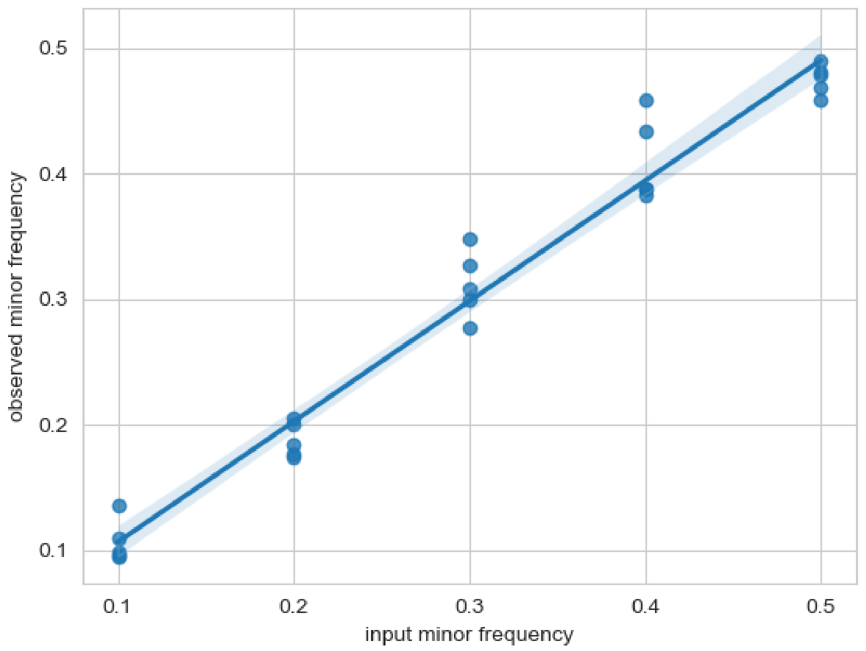



Huang, W., L. Li, J. R. Myers, and G. T. Marth. 2012. 'ART: a next-generation sequencing read simulator', *Bioinformatics*, 28: 593-4.
